# Automated whole-organ histological imaging assisted with ultraviolet-excited sectioning tomography and deep learning

**DOI:** 10.1101/2023.04.22.537905

**Authors:** Lei Kang, Wentao Yu, Yan Zhang, Terence T. W. Wong

## Abstract

Three-dimensional (3D) histopathology involves the microscopic examination of a specimen, which plays a vital role in studying tissue’s 3D structures and the signs of diseases. However, acquiring high-quality histological images of a whole organ is extremely time-consuming (e.g., several weeks) and laborious, as the organ has to be sectioned into hundreds or thousands of slices for imaging. Besides, the acquired images are required to undergo a complicated image registration process for 3D reconstruction. Here, by incorporating a recently developed vibratome-assisted block-face imaging technique with deep learning, we developed a pipeline termed HistoTRUST that can rapidly and automatically generate subcellular whole organ’s virtual hematoxylin and eosin (H&E) stained histological images which can be reconstructed into 3D by simple image stacking (i.e., without registration). The performance and robustness of HistoTRUST have been successfully validated by imaging all vital mouse organs (brain, liver, kidney, heart, lung, and spleen) within 1–3 days depending on the size. The generated 3D dataset has the same color tune as the traditional H&E stained histological images. Therefore, the virtual H&E stained images can be directly analyzed by pathologists. HistoTRUST has a high potential to serve as a new standard in providing 3D histology for research or clinical applications.

## Introduction

Histological imaging of organ slices can provide detailed information about cell morphology and tissue structures, and therefore, it has been widely used in cancer diagnosis by pathologists^1^. Histological images of a few tissue sections are insufficient to reveal the entire organ’s three-dimensional (3D) structural information or internal spatial correspondence. To evaluate the tumor invasiveness or embryonic development^2,3^, 3D histological information with a high axial resolution is needed, which, however, would require hundreds or even thousands of serial tissue slices to be sectioned and stained. The whole procedure can be highly time-consuming (e.g., several weeks) and labor-intensive since the sample has to be preprocessed, sliced, stained, and imaged using different machines with multiple laborious steps (as shown in Fig. 1, marked with a red dashed box). In this circumstance, taking care of hundreds of sections costs several weeks, involves tremendous human labor, and consumes lots of reagents, thus greatly hindering the routine use of 3D histopathology. In addition, as the relative position of each section is lost after sectioning, a complicated registration algorithm has to be implemented to align the hundreds of histological images for 3D reconstruction^4^, not to mention that some of the sections could be ruptured during processing as the thin slices are fragile.

**Fig. 1.**
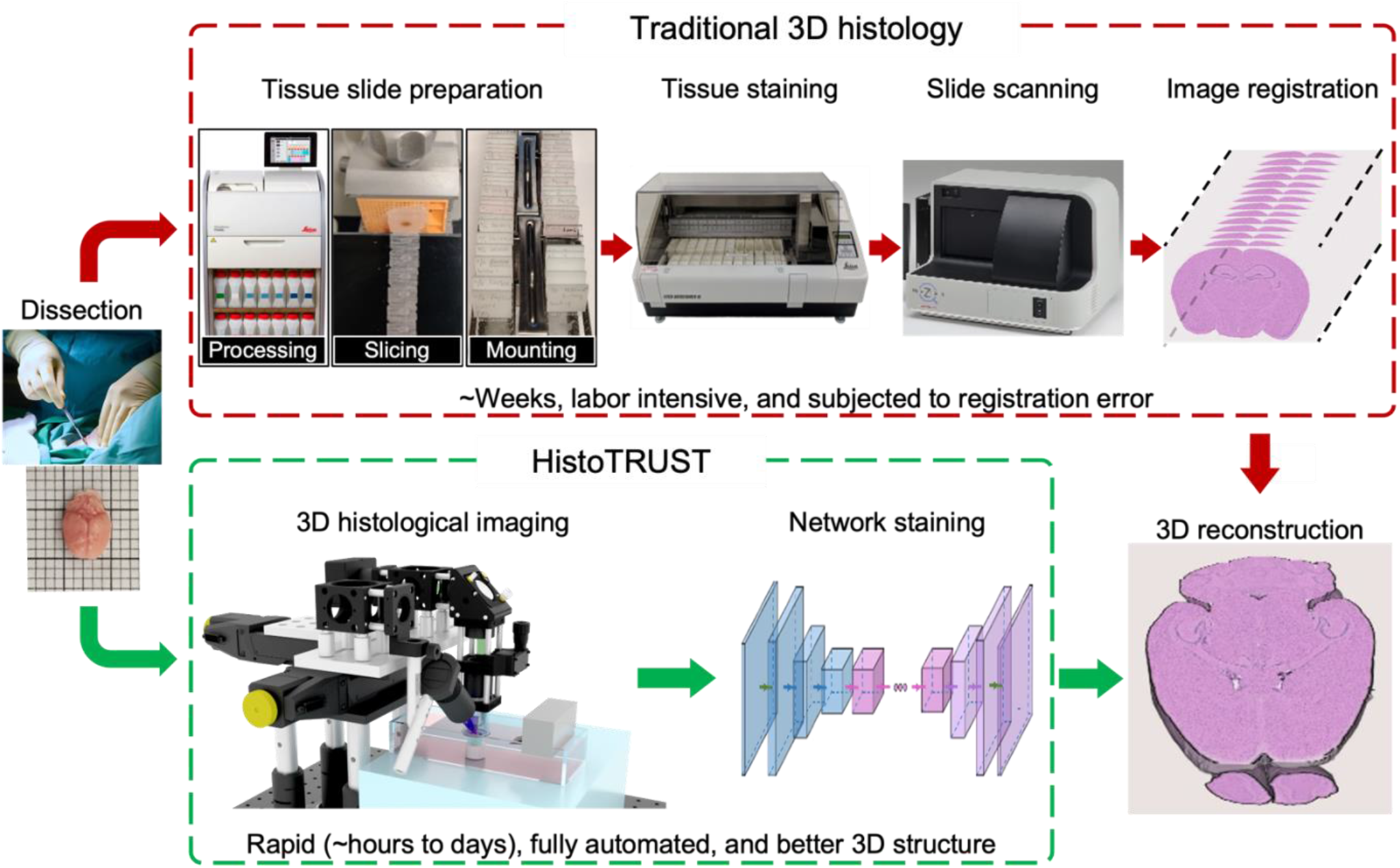
Schematic of the workflow to obtain 3D histological images of whole organs with traditional histopathology and the proposed HistoTRUST. The top pathway shows the conventional approach to obtain 3D H&E stained histological images, which involves lengthy and laborious steps such as tissue slide preparation (e.g., processing, slicing, and mounting), tissue staining, slide scanning, and image registration. The bottom pathway illustrates the proposed HistoTRUST method using automated 3D imaging and deep learning-based virtual staining that can rapidly obtain 3D H&E stained histological images of a whole organ.

Recently, novel imaging methods have been developed to acquire histological images of the whole organ. By integrating photoacoustic microscopy with a microtome, 3D histological images of a mouse brain and kidney have been successfully acquired^5^. However, due to the long scanning time in optical-resolution photoacoustic imaging, it took around 15 days to obtain the 3D histological images of a mouse brain with a sectioning thickness of 200 μm, severely hindering its practical use^6^. Confocal microscopy and light-sheet microscopy also demonstrated their capability to acquire 3D histological images of human brain tissue with the help of modern tissue-clearing techniques, such as CUBIC^7^ and OPTIClear^8^. However, the involved tissue clearing and immunostaining protocols usually take weeks and must be carefully conducted by experienced researchers. In addition, the limited penetration of labeling stains and antibodies into the whole tissue also prevents the further application of the clearing-based methods^9^. Two-photon microscopy can acquire 3D histological images of a mouse brain with serial tissue sectioning in several days, while it is expensive in system cost and especially suitable for organs with inherent fluorescence labels^10^. Our recently proposed microtomy-assisted autofluorescence tomography is another 3D whole-organ histological imaging method, and it is label-free by utilizing the intrinsic contrast of tissue^11^. However, it involves additional paraffin-embedding steps and may lead to tissue shrinkage^11,12^. In contrast to the above sectioning-based methods, magnetic resonance imaging and micro-computed tomography are section-free and can rapidly obtain whole-organ images within one day^13,14^. However, the acquired images have a significant mismatch with conventional histological images in terms of both imaging resolution and molecular contrast, which could hinder the interpretation by pathologists^15^.

Considering the drawbacks of the traditional 3D histological imaging method and the limitations of previously proposed 3D histological imaging modalities, here, we propose a pipeline for the rapid and automated acquisition and generation of virtual hematoxylin and eosin (H&E) stained histological images of whole organs (as shown in Fig. 1, marked with a green dashed box, termed HistoTRUST). HistoTRUST is based on our recently developed high-speed block-face imaging technique named translational rapid ultraviolet-excited sectioning tomography (TRUST), which can acquire serial histological images with a subcellular resolution of whole organs with ultraviolet illumination and vibratome-assisted sectioning^16^. The obtained images will be further transformed into virtual H&E stained histological images by our developed deep-learning neural network. With HistoTRUST, 3D virtual H&E stained histological images of all vital organs in mice, such as the brain, liver, kidney, heart, lung, and spleen, have been acquired in a maximum of three days without any human involvement. In addition, HistoTRUST is compatible with conventional histopathological staining and interpretation techniques. Therefore, it can be directly adopted by pathologists, making it a practical 3D histological imaging tool for research and clinical applications.

## Results

During the imaging process with the TRUST system, we applied two fluorogenic dyes (4’,6-diamidino-2-phenylindole (DAPI) and propidium iodide (PI)) together for double labeling to achieve high color contrast and reveal rich biological information^16^. Once the uppermost layer of the organ has been labeled, the whole section will be raster-scanned by the TRUST imaging system, which is based on the ultraviolet surface excitation for block-face imaging^17^. Then, the imaged surface layer will be sliced off by a vibratome to expose the adjacent layer of the tissue for the next round of staining and image scanning. This process will be repeated automatically until the whole organ has been sectioned and imaged.

We first demonstrate that the TRUST system could provide 2D virtual H&E stained histological images of the organ surface with the assistance of deep learning (See Material and methods). To illustrate that the acquired virtual H&E images can provide similar tissue features as the authentic H&E stained images, the formalin-fixed and paraffin-embedded (FFPE) section adjacent to the imaged organ surface was sectioned and stained with the standard H&E staining protocol for comparison. To illustrate that HistoTRUST can be appliable to different kinds of organs, multiple organs have been imaged, including the brain, liver, kidney, heart, lung, and spleen.

In Fig. 2, the DAPI and PI labeled fluorescence images (called TRUST images) acquired by the TRUST imaging system, corresponding virtual H&E stained histological images transformed from TRUST images, and reference H&E stained histological images are presented. The reference H&E stained histological images were acquired by staining a section (7-μm) adjacent to the previously imaged surface (the TRUST image) following the standard H&E staining protocol, serving as the ground truth for comparison.

**Fig. 2.**
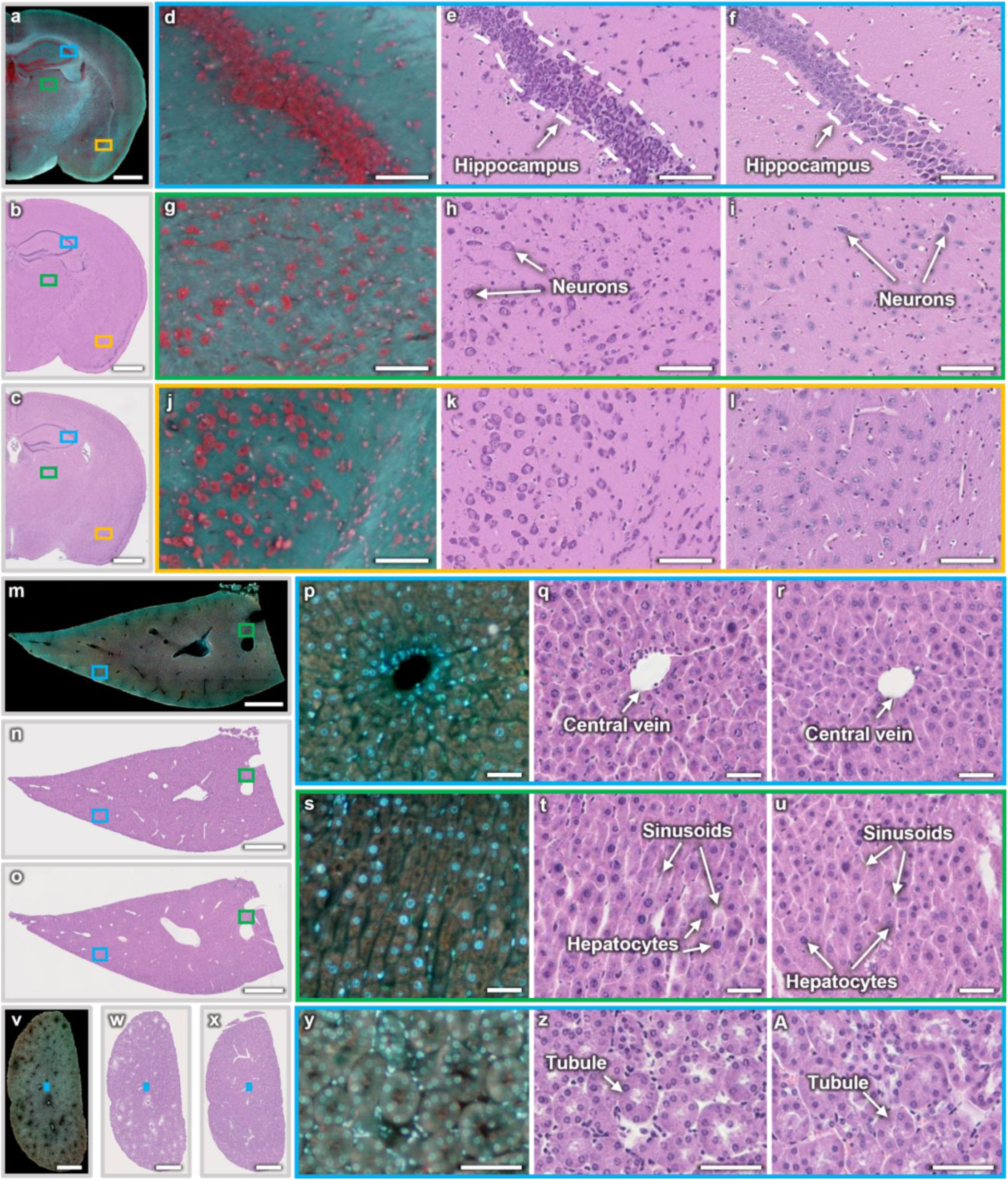
Validation of HistoTRUST on the mouse brain, liver, and kidney. **a**,**b**, TRUST image and its corresponding virtual H&E stained histological image of the brain surface, respectively. **c**, The corresponding reference H&E stained histological image of a section (7-μm) adjacent to the imaged brain surface in a, acquired with traditional histology, serving as the ground truth for comparison. **d–f, g–i, j–l**, Respective zoomed-in TRUST images, virtual H&E images, and reference H&E images showing brain hippocampus, stem, and cortical subplate. **m**,**n**, TRUST image and its corresponding virtual H&E stained histological image of the liver surface, respectively. **o**, The corresponding reference H&E stained histological image of a section (7-μm) adjacent to the imaged liver surface in m. **p–r, s–u**, Respective zoomed-in TRUST images, virtual H&E images, and reference H&E images showing liver cells circulating the central vein and arranging in the radial direction. **v**,**w**, TRUST image and its corresponding virtual H&E stained histological image of the kidney surface, respectively. **x**, The corresponding reference H&E stained histological image of a section (7-μm) adjacent to the imaged kidney surface in v. **y**,**z**,**A**, Zoomed-in TRUST image, virtual H&E image, and reference H&E image showing kidney tubules, respectively. Scale bars: 1 mm (**a–c, m–o, v–x**), 100 μm (**d–f, g–i, j–l**), and 50 μm (**p–r, s–u, y, z, A**).

The TRUST image, its corresponding virtual H&E stained histological image, and the reference H&E stained histological image of a mouse brain surface are shown in Fig. 2a–2c, respectively. To assist the interpretation by pathologists and histopathological analyzing techniques, it is essential to transform the TRUST images into virtual H&E stained histological images. Moreover, three pairs of zoomed-in regions from different functional areas of the brain are present for detailed comparisons, such as the hippocampus structure (Fig. 2e,f), brain stem (Fig. 2h,i), and cortical subplate (Fig. 2k,l). In summary, original TRUST images (Fig. 2a,d,g,j) with different cell densities and morphology were successfully transformed into virtual H&E images (Fig. 2b,e,h,k) in terms of tissue features and color tones.

The TRUST image, its corresponding virtual H&E image, and reference H&E image of a liver are shown in Fig. 2m–2o, respectively. Two regions are zoomed in, showing liver cells that circulate the central vein (Fig. 2p–2r) and arrange in the radial direction (Fig. 2s–2u), respectively. The tissue morphology and cell distributions of the virtual H&E images (Fig. 2q,t) are as authentic as those in the reference H&E images (Fig. 2r,u). As shown in Fig. 2q,r, the virtual and reference H&E stained histological images present a high similarity in terms of the overall style and content. In accordance with our earlier investigation^18^, we also quantitatively evaluated the staining accuracy by comparing the cellular features between the virtual H&E and reference H&E stained images (See Quantitative analysis). Fig. 2v–x are the TRUST image, corresponding virtual H&E image, and reference H&E image of the kidney, respectively. Zoomed-in images (Fig. 2y,z,A) present the important kidney anatomical feature — renal tubules, which are pipe-like structures containing filtered fluid in the kidney^19^. The virtual H&E image provides the same renal tubule structures as the reference H&E image.

More validation results for other organs in mice (e.g., a heart, lung, and spleen) are present in Fig. 3. Fig. 3a–c are the TRUST image, the virtual H&E image, and the corresponding reference H&E image of a heart section, respectively. Two regions are zoomed in to show the cardiac muscle aligned along the longitudinal (Fig. 3d–f) and transversal directions (Fig. 3g–i). Fig. 3j–l are the TRUST image, the virtual H&E image, and the corresponding reference H&E image of the lung section, respectively. Fig. 3m–o and Fig. 3p–r are the corresponding zoomed-in images showing lung alveoli and vessels, respectively. Fig. 3s–u are the TRUST image, the virtual H&E image, and the corresponding reference H&E image of the spleen section, respectively. Fig. 3v–x are corresponding zoomed-in images showing the white pulp structure of the spleen.

**Fig. 3.**
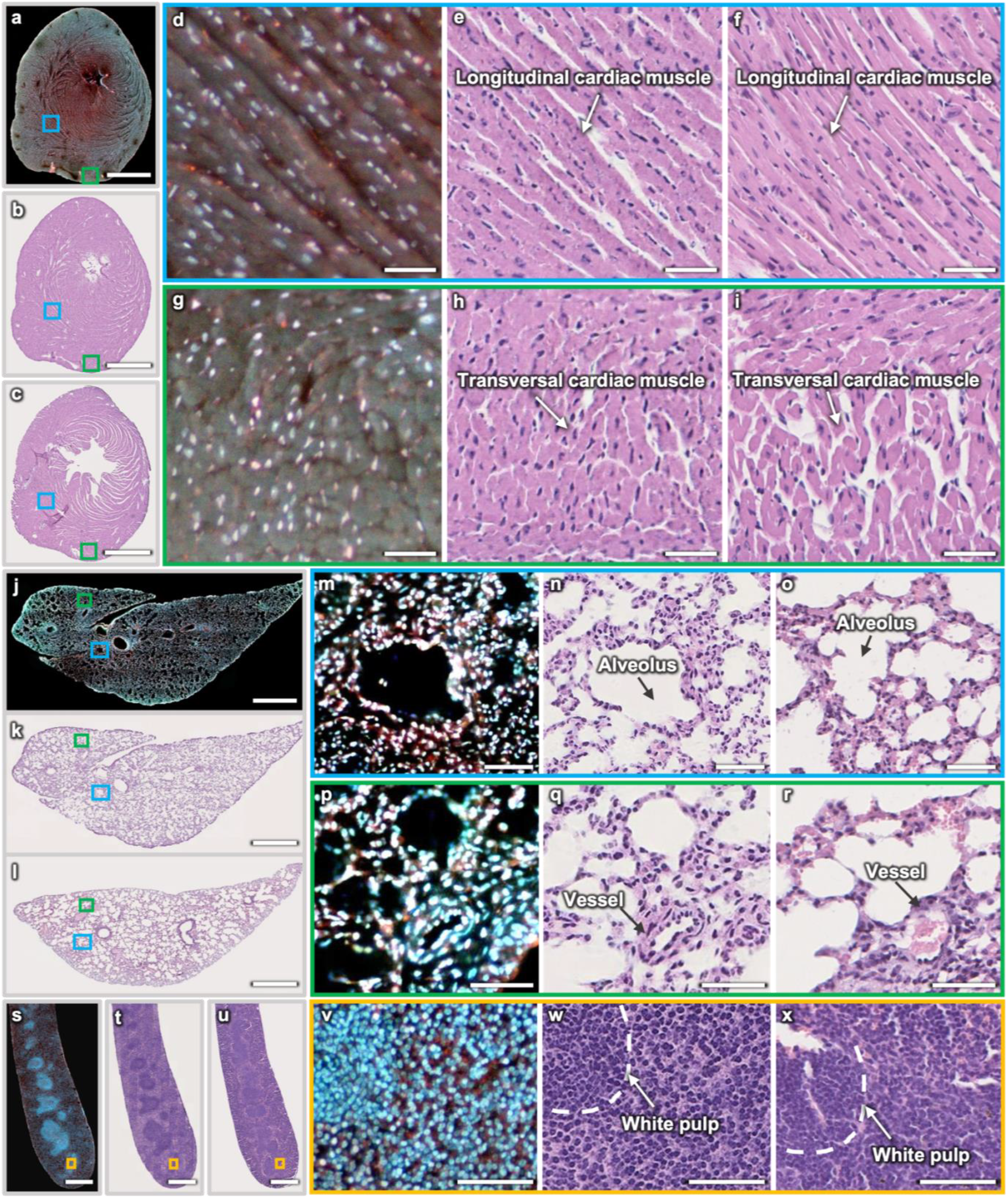
Validation of HistoTRUST on the mouse heart, lung, and spleen. **a**,**b**, TRUST image and its corresponding virtual H&E stained histological image of the mouse heart surface, respectively. **c**, The corresponding reference H&E image of a section (7-μm) close to the imaged heart surface in a, acquired with traditional H&E histology. **d–f, g–i**, Respective zoomed-in TRUST, virtual H&E, and reference H&E images showing cardiac muscle aligned along the longitudinal and transversal directions. **j**,**k**, TRUST image and its corresponding virtual H&E image of the imaged lung surface, respectively. **l**, The corresponding H&E image of a section (7-μm) close to the imaged lung’s surface in j. **m– o, p–r**, Respective zoomed-in TRUST, virtual H&E, and reference H&E images showing lung alveoli and blood vessels. **s**,**t**, TRUST image and its corresponding virtual H&E image of the spleen surface, respectively. **u**, The corresponding reference H&E image of a section (7-μm) next to the imaged spleen surface in s. **v–x**, Zoomed-in TRUST, virtual H&E, and reference H&E images showing the white pulp structure of the spleen, respectively. Scale bars: 1 mm (**a–c, j–l, s– u**) and 50 μm (**d–f, g–i, m–o, p–r, v–x**).

HistoTRUST can provide high-quality 2D virtual H&E stained histological images of various organs (Figs. 2 and 3). The 3D imaging of the entire organ can be reconstructed with hundreds of serial 2D virtual H&E stained histological images. The spatial information of all serial histological images is well known due to the block-face imaging setup and the high-precision translation stages. Therefore, the 3D structures of organs can be easily reconstructed by stacking all acquired 2D images without any image registration. We further present the reconstructed 3D histological images of various mouse organs in Figs. 4 and 5.

**Fig. 4.**
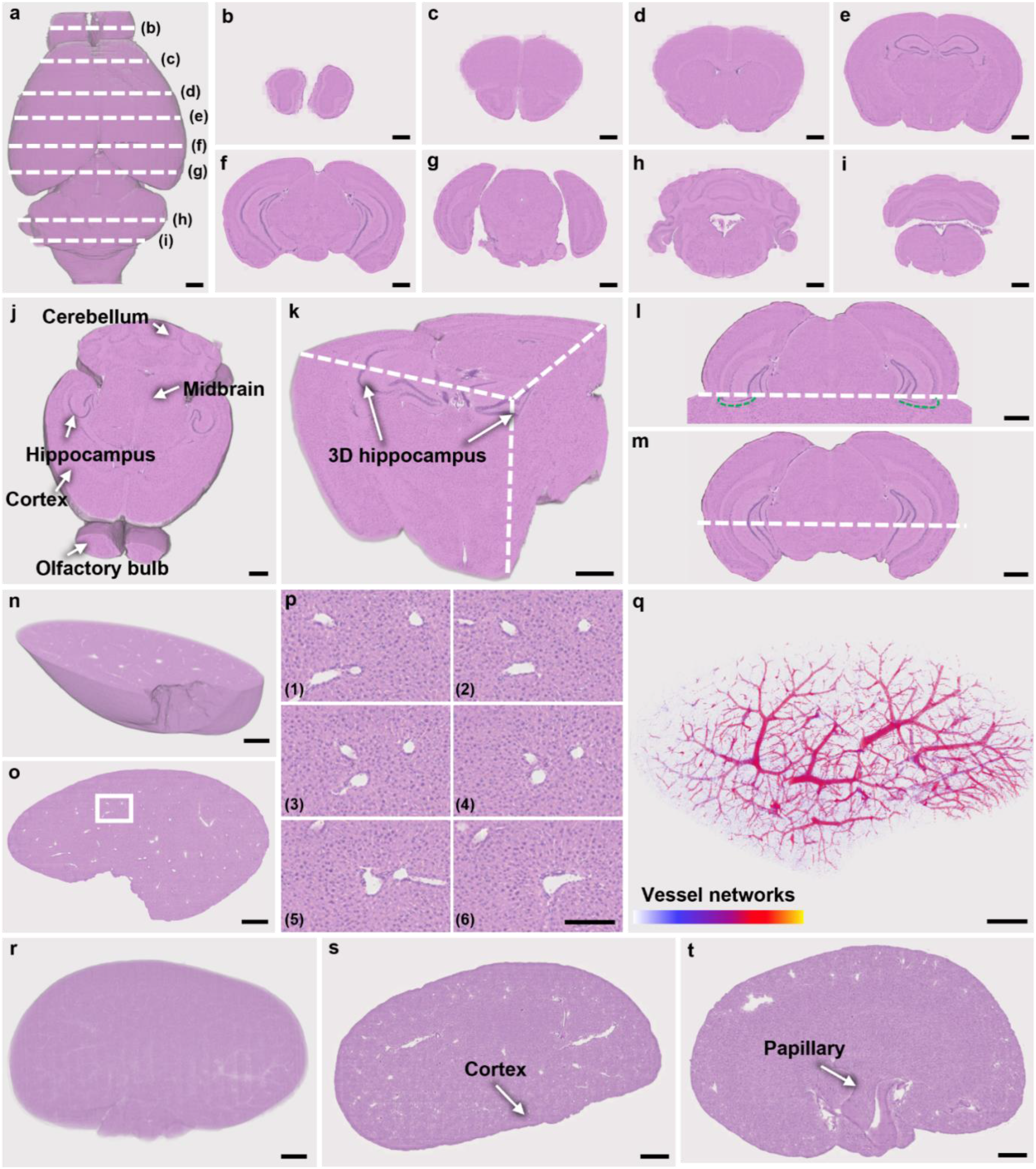
3D volumetric rendering and 3D visualization of anatomical features of a mouse brain, liver, and kidney. **a**, 3D histological image of the whole mouse brain reconstructed with hundreds of serial virtual H&E stained histological images. **b–i**, Virtual H&E images of coronal sections at different depths of the mouse brain. **j–m**, Extracted 3D features of the mouse brain with different perspectives. **n**, A reconstructed 3D histological image of a mouse liver block. **o**, A representative 2D virtual H&E image of the imaged liver block in n. **p**, Serial zoomed-in images of white box region in o. **q**, Vessel networks of the liver block in n extracted by image thresholding. **r**, A reconstructed 3D histological image of a mouse kidney. **s**,**t**, Two virtual H&E images at different depths of the kidney, showing cortex and papillary structures, respectively. Scale bars: 1 mm (**a–o, q–t**) and 50 μm (**p**).

**Fig. 5.**
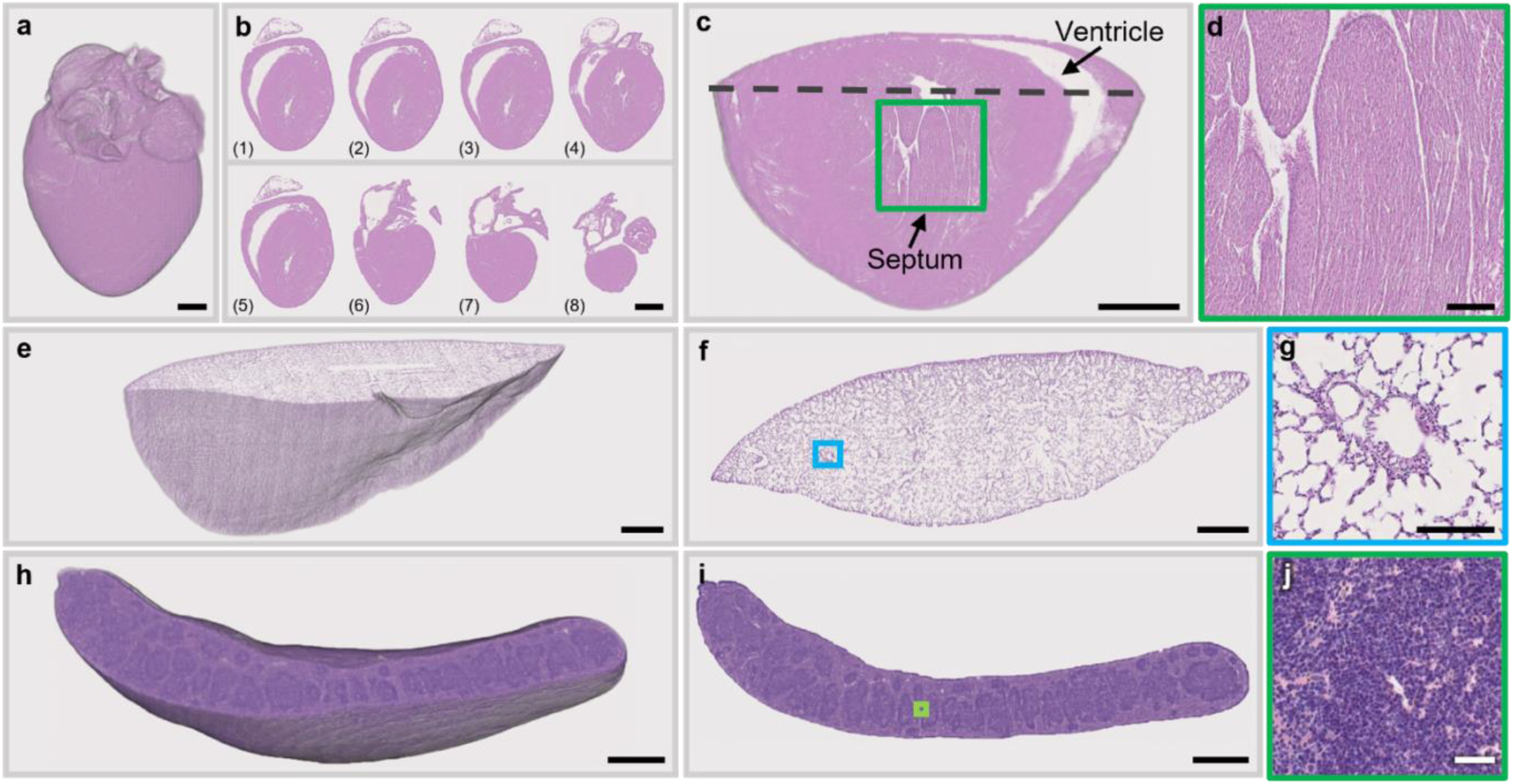
3D volumetric rendering and 3D visualization of anatomical features in the mouse heart, lung, and spleen. **a**, A reconstructed 3D histological image of the mouse heart. The aorta and pulmonary can be observed. **b**, Virtual H&E images showing different sections of the mouse heart. **c**, 3D volumetric rendering showing 3D features of the mouse heart (heart ventricles and interventricular septum). **d**, Zoomed-in 2D H&E image of the green box region in c. **e**, A reconstructed 3D histological image of a mouse lung. **f**, A representative 2D virtual H&E image of the mouse lung. **g**, Zoomed-in 2D H&E image of the blue box region in f, showing lung features (blood vessels and alveoli). **h**, A reconstructed 3D histological image of a mouse spleen. **i**, 2D virtual H&E image of the mouse spleen. **j**, Zoomed-in 2D H&E image of the green box region in i, showing the white pulp structure in the spleen. Scale bars: 2 mm (**b**), 1 mm (**a**,**e**,**f**,**h**,**i**), 500 μm (**c**), 100 μm (**d**), and 50 μm (**g**,**j**).

Figure 4a is the reconstructed 3D virtual H&E histological image of the whole mouse brain with its representative coronal sections presented in Fig. 4b–i, showing major anatomical features, such as the olfactory bulb, cortex, hippocampus, midbrain, and cerebellum^20,21^. The virtual H&E images present a high consistency with the public mouse brain model: Allen mouse brain atlas^22^. 2D virtual H&E images are limited to the planar structure, while more information can be obtained with the reconstructed 3D model. For example, as shown in Fig. 4j, the sagittal view of the mouse brain can also be reconstructed, showing different brain structures simultaneously, such as the cerebellum, hippocampus, cortex, and olfactory bulb. Also, the volumetric rendering (Fig. 4k) reveals the extension of brain features in 3D space (e.g., the hippocampus in 3D). Fig. 4l further shows that the inner hippocampus is spatially connected with the outer hippocampus, which cannot be recognized from the 2D coronal section (Fig. 4m). Fig. 4n presents the reconstructed 3D histological image of a mouse liver block, and Fig. 4o shows one of the 2D virtual H&E liver images used for 3D reconstruction. We can trace the 3D extension of the blood vessels at the region marked with a white box in Fig. 4o, and Fig. 4p (1–6) are serial zoomed-in images of the same area from adjacent layers, showing the gradual fusion of several vascular branches into a single vessel. The vessel networks of the whole imaged liver block can also be extracted after image thresholding, as shown in Fig 4q. Fig. 4r is the reconstructed 3D histological image of a mouse kidney, and Fig. 4s,t show two virtual H&E images at different depths in the kidney, showing cortex and papillary structures^23,24^, respectively.

Figure 5a shows the reconstructed 3D histological image of an entire mouse heart. In addition to the 3D morphology of the heart, the aorta, and pulmonary veins which compose the blood circulation system^25,26^, can also be observed. Several virtual H&E images of heart sections are shown in Fig. 5b, which are consistent with histological images from existing mouse heart atlas^27^. Fig. 5c is a zoomed-in volume extracted from Fig. 5a to show the heart ventricle and interventricular septum in 3D space. The interventricular septum is composed of cardiac muscle and fibrous tissue^28^, which is further illustrated in the zoomed-in 2D histological image (Fig. 5d). Fig. 5e is the reconstructed 3D histological image of a mouse lung with one representative 2D virtual H&E lung image shown in Fig. 5f. Fig. 5g is the zoomed-in image of the blue box region in Fig. 5f, showing lung vessels, alveoli, and individual cells. Fig. 5h is the 3D histological image of a mouse spleen with one representative 2D virtual H&E image shown in Fig. 5i. Fig. 5j is the zoomed-in image of the green box region in Fig. 5i, showing the white pulp structure in the spleen.

## Discussion

HistoTRUST paves the way to rapidly and automatically acquire virtual H&E histological images of entire organs in 3D, which dramatically relieves researchers from the laborious tissue processing, sectioning, and imaging scanning procedures by integrating them into a single and automated 3D imaging acquisition and virtual staining system. HistoTRUST is time-efficient and can realize whole-organ imaging within hours to days, depending on the volume of imaged samples, by removing the common tissue processing steps (e.g., the paraffin embedding or H&E histological staining). HistoTRUST is also compact and cost-efficient because bulky and expensive machines used in conventional 3D histology for tissue processing and staining are unnecessary. Besides, to the best of our knowledge, HistoTRUST is the only 3D histological imaging method that has been verified on all vital organs in mice (e.g., brain, liver, kidney, heart, lung, and spleen), which shows its wide applicability. Moreover, unlike some other 3D histological imaging modalities where the output images are different from the H&E stained images in terms of color tones and molecular features, the acquired virtual H&E stained histological images from HistoTRUST can be directly used by pathologists for diagnosis due to its high similarity with H&E stained images, and therefore, are completely compatible with conventional histopathological interpretation techniques. Overall, the proposed HistoTRUST is a reliable and powerful tool for 3D histological imaging in terms of speed, equipment and labor cost, applicability, and compatibility, thus promoting the use of 3D histopathological analysis in research or clinical applications.

HistoTRUST is at its early developing stage and can be further improved in the following aspects. Limited by the sectioning performance of the vibratome model that is currently used, the entire organ was imaged at ∼50-μm interval, which could be reduced to ∼4 μm with the latest vibratome models (e.g., VF-300-0Z, Precisionary Instruments Inc.), thus offering better continuity and better axial resolution for the reconstructed 3D histological image. In addition to improving its imaging ability, more experiments can be conducted to explore its potential application in the medical and life science domains. For example, tumor development in 3D space can be analyzed by acquiring 3D histological images of different tissue samples with tumors. With these further development, HistoTRUST could be a practical and universal tool for whole-organ 3D histopathological analysis in research or clinical settings.

## Material and methods

### Sample preparation

Once mice (C57BL/6) were sacrificed, the internal organs were harvested immediately and rinsed with phosphate-buffered saline solution for a minute. Next, the organs were submerged under 10% neutral-buffered formalin at room temperature for 24 hours for fixation, and then they were directly imaged by the HistoTRUST system. All animal experiments were conducted in conformity with a laboratory animal protocol approved by the Health, Safety, and Environment Office of the Hong Kong University of Science and Technology (HKUST).

We also acquired the H&E stained histological image for each type of organ with the traditional FFPE H&E staining protocol for training the respective neural networks that can transfer the acquired TRUST images to virtual H&E stained histological images. After fixation, the extracted organ undertook a series of operations, including dehydration, clearing, and paraffin infiltration within a Revos Tissue Processor (Thermo Fisher Scientific Inc.), consuming nearly 13 hours. Subsequently, the organ was embedded into a paraffin block using a HistoStar Embedding Workstation (Thermo Fisher Scientific Inc.) and sectioned into 7-μm thin slices by a microtome. The tissue slices were mounted on glass slides, dried in an incubator for one hour, and then deparaffinized with xylene. After deparaffinization, the tissue slices were stained with H&E and imaged by a whole-slide imaging machine (Nanozoomer-SQ C13140, Hamamatsu Photonics K.K.) with a 20×/numerical aperture 0.75 objective lens (CFIPlanApo Lambda 20×, Nikon Corp.) to obtain H&E stained histological images. The H&E stained histological image acquisition process only needs to be done once for network training.

### Image acquisition

As shown in Fig. 6a, the agarose-embedded organ was submerged under the mixed fluorescent dyes (for a brain: 5 μg/ml DAPI and 5 μg/ml PI; for other organs: 5 μg/ml DAPI and 2.5 μg/ml PI) contained in the water tank of the vibratome to label cell nucleus of the organ’s topmost layer within ∼2.5 minutes. Then, the fully stained surface was illuminated with an ultraviolet light-emitting diode (UV-LED) (∼285 nm, M285L5, Thorlabs Inc.) to excite fluorescence signals which were collected by a 10× objective lens (RMS10X, Olympus Corp.), focused by an infinity-corrected tube lens (ACA254-200-A, Thorlabs Inc.), and finally recorded by a color camera (DS-Fi3, Nikon Corp.). With a two-axis motor stage (L-509.10SD00, Physik Instruments (PI) GmbH & Co.KG), the fluorescence signals of the entire surface layer can be acquired by raster scanning and image stitching. Finally, the previously imaged layer will be removed by the blade of the vibratome (VF-700-0Z, Precisionary Instruments Inc.) with a thickness of ∼50 μm to expose an adjacent layer for the next round of tissue staining, imaging, and sectioning. The process would be automatically repeated hundreds of times until the entire organ has been imaged.

**Fig. 6.**
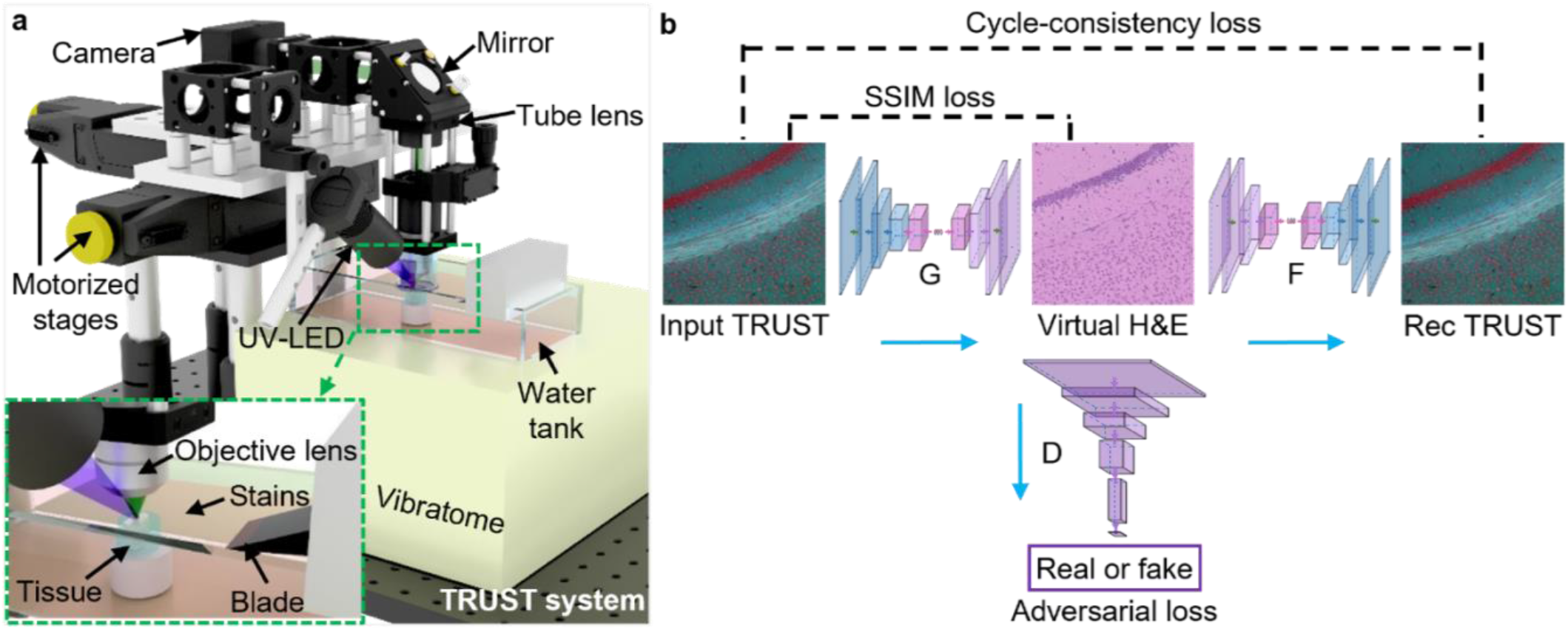
Illustration of the HistoTRUST imaging system and CycleGAN virtual staining network. **a**, The schematic of the HistoTRUST imaging system that generates serial histological images (TRUST images) of whole organs. **b**, The detailed workflow of transforming TRUST images into virtual H&E stained histological images using CycleGAN. The generator G transforms the TRUST images to the corresponding virtual H&E images, and then the discriminator D distinguishes the generated virtual H&E images from the authentic H&E stained images. Generator G tries to generate virtual H&E images that D cannot discriminate. The SSIM loss is calculated between the input TRUST images and the generated virtual H&E images. A generator F further transforms the virtual H&E images back to TRUST images (Rec TRUST), which will be used to calculate the cycle-consistency loss with the original TRUST images.

The whole-organ imaging procedure is fully automated and controlled by a customized control system based on LabVIEW (National Instruments Corp.). The customized software provides a user-friendly interface to adjust key imaging parameters such as the step size and travel range of the motorized stages, which also ensures the correct sequence of vibratome sectioning, staining, and whole surface imaging. To synchronize the sequence of motor scanning and camera imaging, we customized the camera imaging software with Visual Studio to enable the external trigger function of the DS-Fi3 color camera. Finally, to control the operation of the vibratome, which has no external triggering interface of its own, we customized a simple electrically controllable mechanical device based on stepper motors to push the buttons on the vibratome’s control panel.

### Deep learning-based virtual staining neural network

The HistoTRUST imaging system can rapidly acquire histological TRUST images of the whole organ. Then, a virtual staining network can instantly transform the TRUST images into virtual H&E stained histological images to assist the image interpretation by pathologists, promoting its use in research or clinical applications.

We adopt the cycle-consistent generative adversarial network (CycleGAN) architecture^29^ for virtual staining. The CycleGAN consists of two types of networks: the generators and the discriminator. The generator tries to produce images that the discriminator cannot distinguish^30^ during the network training. As shown in Fig. 6b, generator G transforms the TRUST images to virtual H&E images, and then discriminator D distinguishes the generated virtual H&E images from the authentic H&E stained images. Once generator G can produce virtual H&E images that discriminator D cannot distinguish, it acquires the ability of virtual H&E staining digitally. The generated virtual H&E image is also transformed back to a TRUST image (Rec TRUST) to implement cycle-consistency loss, which helps to stabilize the network training. The generator and discriminator are fully convolutional neural networks, and the detailed layer compositions are shown in supplementary Fig. S1. We made two improvements to the original generator from CycleGAN^29^. First, we reduced the kernel size of the first convolutional layer from 7 × 7 to 3 × 3, which helps to focus on small features and reduce the computational cost. Second, the transposed convolutional layer is substituted with the pixel shuffle layer^31^, which can remove the random checkbox artifacts in the original CycleGAN (supplementary Fig. S2).

The CycleGAN contains three types of losses: adversarial loss^32^, cycle-consistency loss^29^, and identity loss. To better preserve the original structure information in TRUST images, the structural similarity index loss (SSIM)^33^ is added between the TRUST and virtual H&E images. The objective for SSIM can be expressed as:

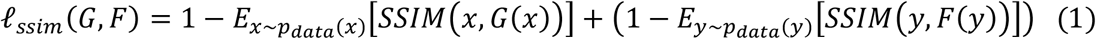

where *x* and *y* represent images from the TRUST and H&E domains, respectively.

### Implementation details Image pre-processing

The acquired TRUST images have three channels (RGB), and the R channel was selected for training as it contains information about the cell nucleus and is less affected by the autofluorescence signals excited by the UV light. The TRUST image was then cropped into small tiles (256 × 256 pixels) by a sliding window with a moving step of 150 pixels. For the H&E image, it was firstly resized to match the scale size of nuclei in the TRUST image before undergoing the same image cropping algorithm. Since CycleGAN is an unsupervised learning method that can be trained on unpaired data, the TRUST and H&E images used for training can be unregistered, thus bypassing the complex image registration process and also easing the acquisition of training datasets compared with previously reported supervised-based virtual staining methods^34,35^.

### Network training

Six CycleGAN neural networks have been trained respectively for six different types of organs. Each CycleGAN network was trained on a dataset composed of 1,600 TRUST and 1,600 H&E patches. The TRUST and H&E images used for training can be unregistered because of the unpaired training superiority of CycleGAN. Therefore, typical FFPE thin slices stained with standard H&E staining protocol can be directly used to ease the acquisition of training datasets. The CycleGAN network was implemented with PyTorch (version 1.0.1), trained and tested on a desktop computer running Ubuntu (18.04.2 LTS) operation system, with Core i7-9700K CPU@ 3.6GHz, 64GB RAM, and an NVIDIA Titan RTX GPU. Each CycleGAN network was trained for 100 epochs, taking around 8 hours.

The cycle-consistency loss serves as an indicator to terminate the training. As shown in Fig. 7a, after 100 epochs of training, the loss converged to a low level, and the trained networks can output high-quality virtual H&E stained images.

**Fig. 7.**
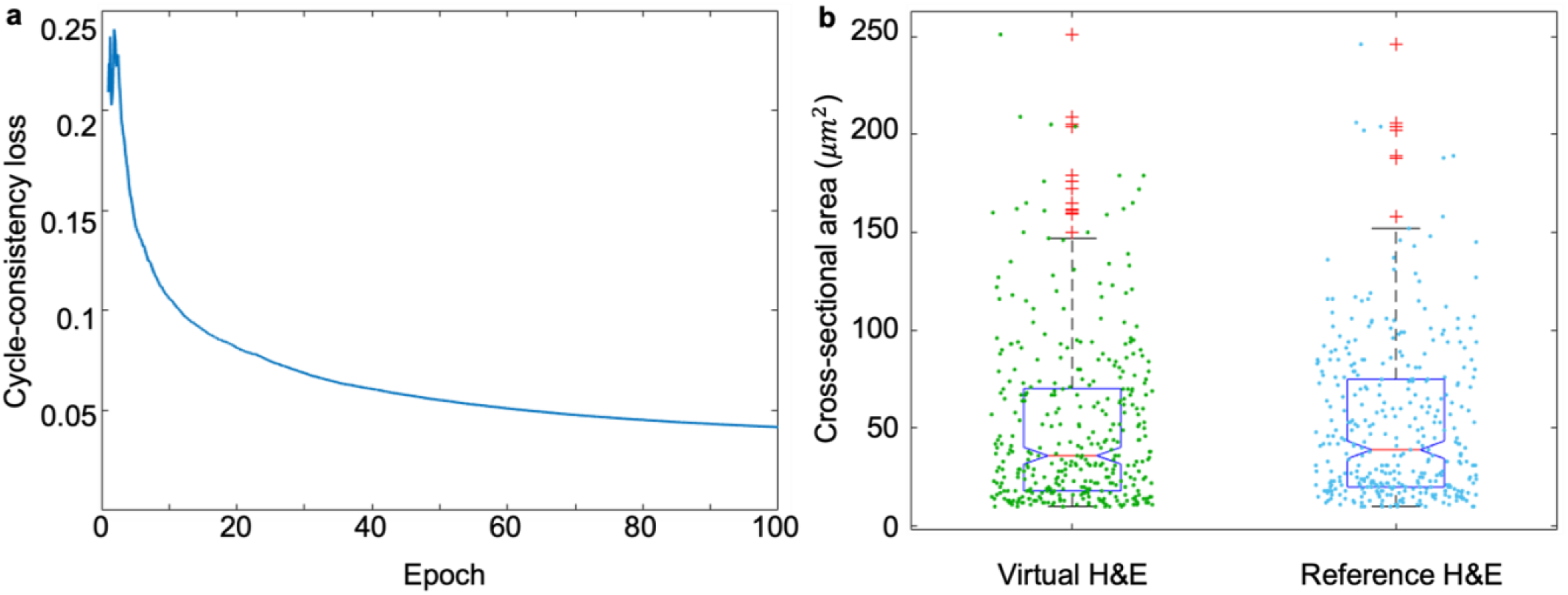
CycleGAN training loss and quantitative analysis of virtual H&E stained histological images. **a**, The cycle**-** consistency loss of TRUST images in the CycleGAN training. **b**, Respective cross-sectional area distributions of nuclei in virtual H&E and reference H&E images.

### Quantitative analysis

To evaluate the staining accuracy of our method using HistoTRUST, we quantitatively compared the cellular features between the virtual H&E and reference H&E stained images. A quantitative cellularity analysis about the number of cells (cell counting) and their cross-sectional areas are made on the same virtual H&E and reference H&E images (Fig. 2q,r). The number of cells and the average nuclear cross-sectional area are listed in Table 1, presenting a good match. The cross-sectional area distributions are further plotted in Fig. 7b. The Wilcoxon rank-sum test^36^ is performed for the two distributions, with a null hypothesis that the two datasets come from the same distribution and a significance level of 0.05. The resulting p-value was 0.42, indicating no significant difference between the two nuclear cross-sectional area distributions of virtual and reference H&E images.

**Table 1.** Counts (i.e., number of cells) and the average nuclear cross-sectional area of virtual H&E and reference H&E images (Fig. 2q,r)

| | Counts | Average nuclear cross-sectional area ($\mu\text{m}^2$) |
| --- | --- | --- |
| Virtual-H&E | 369 | 51.02 |
| Reference-H&E | 365 | 51.13 |

## Acknowledgments

The Translational and Advanced Bioimaging Laboratory in HKUST acknowledges the support of the Research Grants Council of the Hong Kong Special Administrative Region (16208620 and 16206522); The Hong Kong University of Science and Technology startup grant (R9421).

## Conflicts of interest

T.T.W.W. has a financial interest in PhoMedics Limited, which did not support this work. W.Y., L.K., Y.Z., and T.T.W.W. have applied for a patent (US Provisional Patent Application No.: 63/254,546) related to the work reported in this manuscript. All authors declare no competing financial interests.

## Author Contributions

L.K. and W.Y. contributed equally to the work. L.K., W.Y., and T.T.W.W. conceived of the study. L.K. built up the control system and pipeline of HistoTRUST, applied the CycleGAN networks for virtual histological staining, and analyzed the results. W.Y. designed and built the optical imaging system of TRUST and performed experiments to acquire the data. Y.Z. gave suggestions on the optical setup. L.K., W.Y., and T.T.W.W. wrote the manuscript. T.T.W.W. supervised the whole study.

## Data availability

The authors declare that all data supporting the findings of this study are available within the paper and its supplementary information.

