## Supplementary Figures 1-2 for "Automated whole-organ histological imaging assisted with ultraviolet-excited sectioning tomography and deep learning"

**The Supplementary Figures:**

Supplementary Fig. 1 | Illustration of the layer compositions in the generator and discriminator derived from the original CycleGAN.

Supplementary Fig. 2 | Enhancing virtual staining performance by substituting the transpose convolutional layer with the pixel shuffle layer.

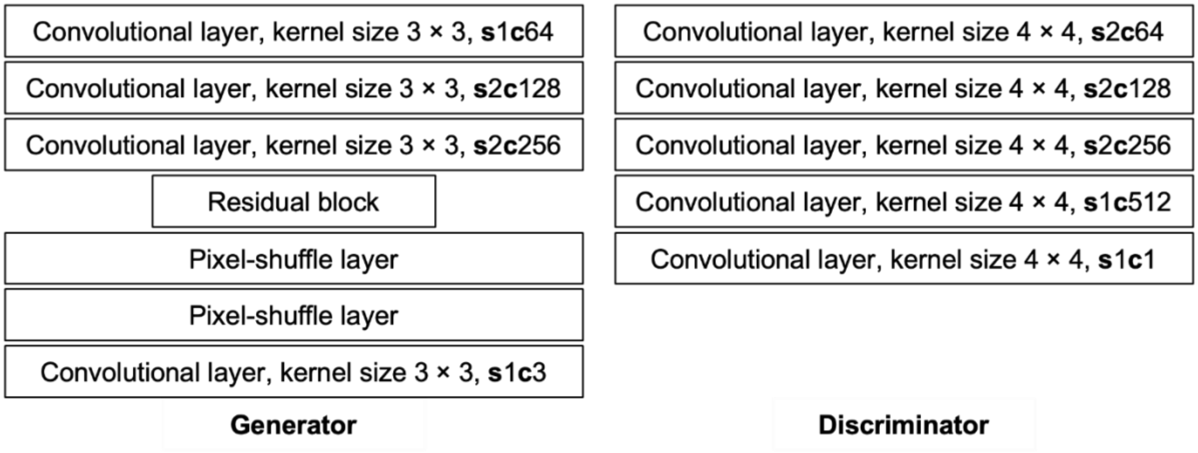


**Supplementary Fig. 1 Illustration of the layer compositions in the generator and discriminator derived from the original CycleGAN.** Left: the composition of layers in the generator. Right: the composition of layers in the discriminator. **s**: stride length; **c**: channel number.

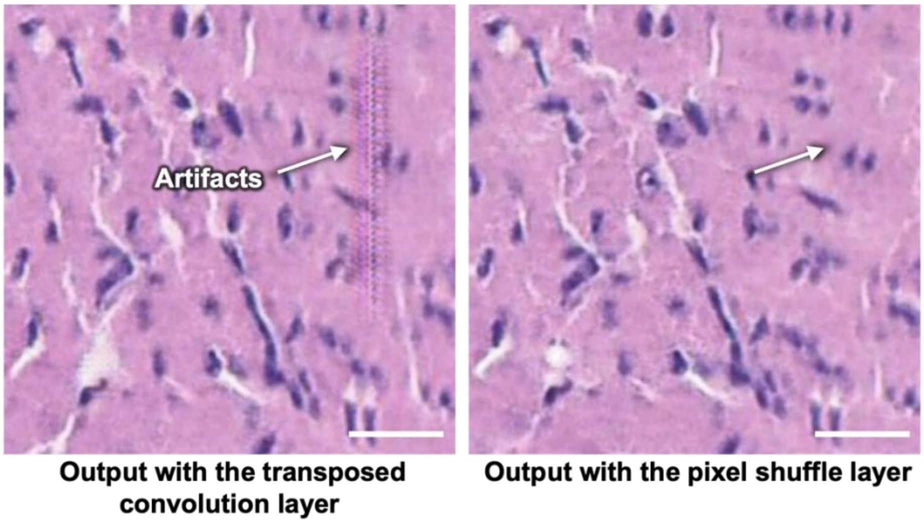


**Supplementary Fig. 2 Enhancing virtual staining performance by substituting the transpose convolutional layer with the pixel shuffle layer.** Left: virtual H&E stained histological image produced by the generator with the transposed convolutional layer, showing random checkbox artifacts. Right: virtual H&E stained histological image produced by the generator with the pixel shuffle layer, showing no artifacts. Scale bars: 50 μm.
